# Conformational control of translation termination on the 70S ribosome

**DOI:** 10.1101/226837

**Authors:** Egor Svidritskiy, Andrei A. Korostelev

## Abstract

Translation termination ensures proper lengths of cellular proteins. During termination, release factor (RF) recognizes a stop codon and catalyzes peptide release. Conformational changes in RF are thought to underlie accurate translation termination. If true, the release factor should bind the A-site codon in inactive (compact) conformation(s), but structural studies of ribosome termination complexes have only captured RFs in an extended, active conformation. Here, we identify a hyper-accurate RF1 variant, and present crystal structures of 70S termination complexes that suggest a structural pathway for RF1 activation. In the presence of blasticidin S, the catalytic domain of RF1 is removed from the peptidyl-transferase center, whereas the codon-recognition domain is fully engaged in stop-codon recognition in the decoding center. RF1 codon recognition induces decoding-center rearrangements that precede accommodation of the catalytic domain. Our findings suggest how structural dynamics of RF1 and the ribosome coordinate stop-codon recognition with peptide release, ensuring accurate translation termination.

## Introduction

Accurate translation termination prevents the premature release and accumulation of potentially toxic polypeptides. At the end of translation, protein release factor (RF) binds the stop codon of an mRNA in the ribosomal A (<u>*a*</u>minoacyl-tRNA) site and catalyzes the hydrolysis of peptidyl-tRNA, releasing the complete polypeptide. In eubacteria, RF1 recognizes the UAA and UAG stop codons, whereas RF2 recognizes the UAA and UGA stop codons (reviewed in refs (1-5)). High fidelity of translation termination by RF1 and RF2, with error rates in the range of 10^−3^ to 10^−6^ (6), equals to or exceeds that of aminoacyl-tRNA (aa-tRNA) incorporation by elongation factor EF-Tu. Accurate decoding of aa-tRNA•EF-Tu requires GTP hydrolysis (7, 8), which separates accommodation of cognate tRNA and dissociation of non-cognate tRNA (9). By contrast, accurate termination by RF1 and RF2 does not employ GTP (6, 10).

One model to explain the accuracy of translation termination is that stop-codon recognition triggers a large-scale conformational rearrangement that activates release factor (11-14). Many structural studies of 70S ribosome termination complexes formed with RF1 or RF2 (11, 12, (15-21)) have revealed an active, i.e. extended, conformation of release factor. In this conformation, the codon-recognition domain II of RF is bound to the stop codon in the 30S A site. Here, the conserved ribosomal decoding-center nucleotides A1492, A1493 (16S rRNA) and A1913 (23S rRNA) adopt a conformation characteristic of 70S termination complexes (18, 19). The catalytic domain III of RF contains a long helix α7 that bridges the ribosomal subunits. The conserved catalytic GGQ motif at the tip of domain III binds next to the CCA 3’-end of the P-site tRNA in the peptidyl-transferase center of the 50S subunit to mediate peptidyl-tRNA hydrolysis (11, 15, (17-20)). Recent biochemical and biophysical data suggest that RF initially samples the ribosomal A site in a conformation that is different from the fully extended one (13, 14). Crystal structures of free RF1 and RF2 have revealed a compact conformation (22-24), but this conformation is sterically incompatible with a fully accommodated codon-recognition domain (18). In solution, free release factors might sample compact or partially extended conformations that are compatible with initial binding in the ribosomal A site (25, 26). However, so far only the fully extended conformation has been captured on the terminating ribosome by electron cryo-microscopy (cryo-EM) or X-ray crystallography. Recently, ribosome rescue complexes formed with alternative rescue factor ArfA captured a compact conformation of RF2 (27, 28). The rescue complex, however, depends on recognition of a truncated (non-stop) mRNA by ArfA, so these structures do not provide insight into how stop-codon recognition mediates peptide release. Thus, structures of RF intermediates that support conformational control of translation termination on stop codons remain elusive.

If conformational rearrangement of release factor underlies termination fidelity, then release factor should initially bind the ribosome with its codon-recognition domain in the decoding center, while the catalytic domain is not yet activated to present the GGQ motif in the peptidyl-transferase center. Therefore, perturbation of RF1 dynamics or perturbation of the catalytic domain binding to the 50S subunit may change the efficiency of termination and yield pre-activation-like states. To test this hypothesis, we first identified a hyper-accurate variant of RF1 (Δ302-304 RF1; RF1_ha_), whose discrimination against sense codons exceeds that of wild-type RF1. We then employed RF1_ha_ to capture distinct termination complexes by X-ray crystallography. To induce a pre-activation-like state, we used the termination inhibitor blasticidin S (BlaS). BlaS intercalates within the CCA end of the P-site tRNA in the peptidyl-transferase center, resulting in partial occlusion of the A site on the 50S subunit (29). We reasoned that BlaS might prevent the catalytic domain of RF1_ha_ from stably binding the 50S subunit, providing insights into early steps of termination.

## Results and Discussion

### Δ302-304 RF1 (RF1_ha_) demonstrates enhanced discrimination against sense codons

In release factors, the codon-recognition superdomain (domains II and IV) is connected to catalytic domain III via a non-conserved loop, termed the switch loop (aa 291-307 of *E. coli* RF1; (11, 18)). In structures of 70S termination complexes (11, 15, 17, 18, 20, 21), the switch loop interacts with decoding-center nucleotides A1492 and A1493 of 16S rRNA and helix 69 (capped with A1913) of 23S rRNA (11, 18). These nucleotides adopt termination-specific conformations, which differ from those observed with a vacant A site (30). Presently, it is unclear whether the switch loop contributes to the decoding-center conformational change that is compatible with extended activated RF in the 70S complex (1). The switch loop does not bind the stop codon in 70S structures, consistent with our previous finding that RF1 variants with mutations in the switch loop retain peptide release activity (18). When combined with a deletion of helix 69, however, RF1 with a shortened switch loop (Δ302-304) had a substantially slower rate-limiting step in peptide release (18). Because the switch loop connects the two functional domains of RF, we hypothesized that the rate-limiting step could be a conformational change that directs domain III into the peptidyl-transferase center (18). If the Δ302-304 RF1 variant slows a critical conformational change, we now asked whether the Δ302-304 RF1 variant might be more discriminatory than RF1_wt_ against sense codons.

To test the above hypothesis, we used an established *in vitro* kinetic assay (19) that monitors the release of formyl-methionine from the 70S ribosome bound with [^35^S]-fMet-tRNA^fMet^ in the P site (Fig. 1A). We compared the release kinetics of wild-type RF1 and Δ302-304 RF1 from the UAA stop codon or the sense codons CAA, UGG, UAU (Table 1) - i.e., *near-stop* codons that differ from UAA or UAG stop codons in the first, second, or third nucleotide position. Both wild-type and mutant RF1 induced fast release on the UAA codon (Fig. 1B, 1C). The release efficiency (k_cat_/K_M_) of RF1_wt_ on near-stop codons was readily detectable but reduced by ~100-to 1000-fold (Fig. 1B), consistent with reported observations (6). By contrast, the activity of Δ302-304 RF1 on near-stop codons was virtually undetectable (Fig. 1C). We estimate the specificity - defined as (k_cat_/K_M_)^stop^ / (k_cat_/K_M_)^sense^ - of Δ302-304 RF1 to be ~10-to 100-fold higher than that of RF1wt (Fig. 1D, Table 1). Thus, destabilizing the switch-loop interactions with the 30S decoding center dramatically increased the accuracy of stop-codon recognition.

**Figure 1.**
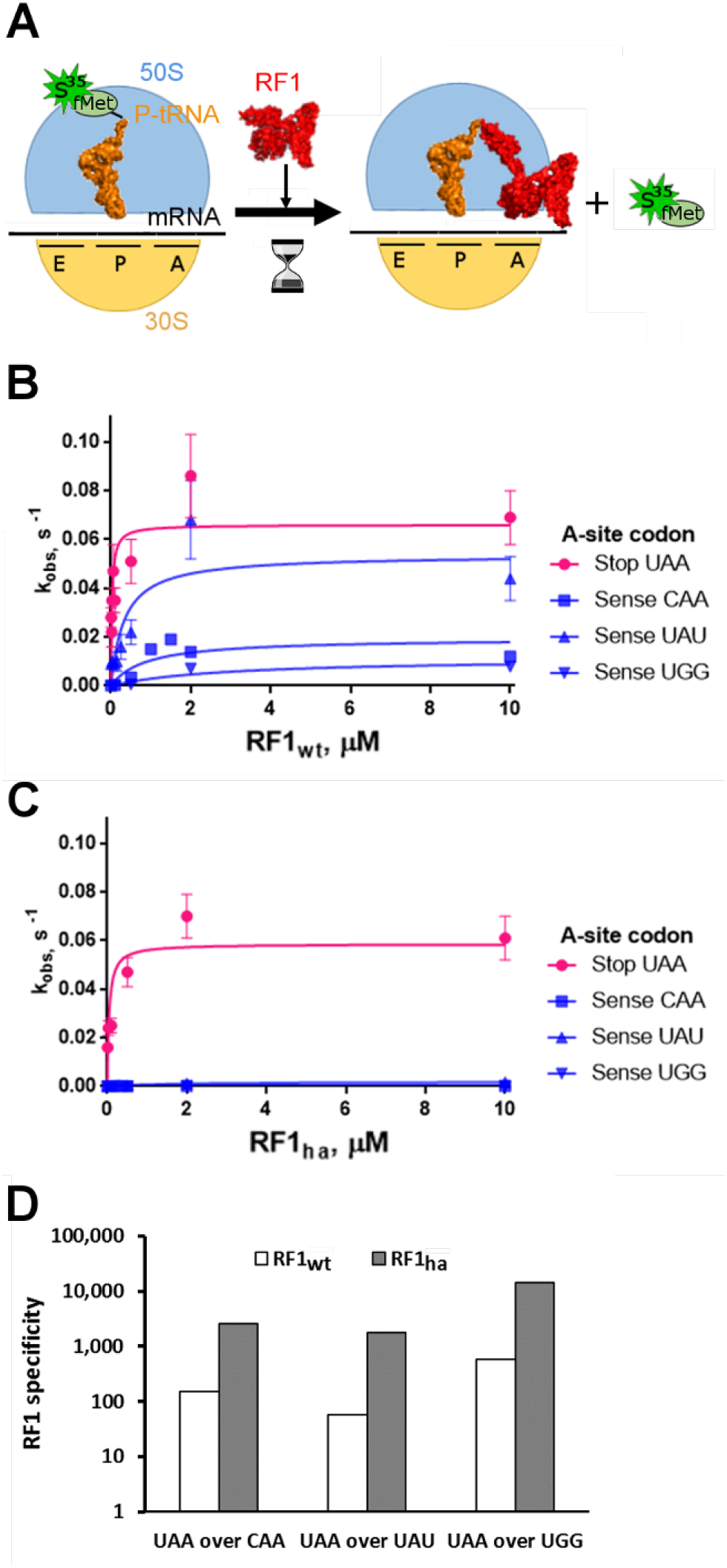
Comparison of termination efficiencies and specificities of *E. coli* RF1_wt_ and Δ302-304 RF1 (RF1_ha_) on 70S ribosomes with a stop or sense codon in the A site. (*A*) Schematic of the experiment to study the kinetics of formyl-methionine release by RF1 from the 70S ribosome bound with fMet-tRNA^fMet^ (see Methods). (*B*, *C*) Michaelis – Menten curves for RF1_wt_ (B) and Δ302-304 RF1 (RF1_ha_) (C); k_obs_ denotes the observed rate constants; error bars represent standard errors obtained from duplicate experiments. (*D*) Specificities, (k_cat_/K_M_)^stop^ / (k_cat_/K_M_)^sense^, of release by RF1_wt_ and Δ302-304 RF1 (RF1_ha_) from the pre-termination ribosome complex (with the UAA stop codon) vs. the elongation complex (with near-stop sense codons), shown on a logarithmic scale.

**Table 1.** Kinetic measurements for fMet release mediated by RF1_wt_ and RF1_ha_.

| **A-site Codon** | **UAA** | **CAA** | **UAU** | **UGG** |
| --- | --- | --- | --- | --- |
| **k<sub>cat</sub> (wt), s<sup>-1</sup>** | $(7.0 \pm 1.0) \cdot 10^{-2}$ | $(1.6 \pm 0.4) \cdot 10^{-2}$ | $(5.1 \pm 1.6) \cdot 10^{-2}$ | $(1.1 \pm 0.4) \cdot 10^{-2}$ |
| **k<sub>cat</sub> (ha), s<sup>-1</sup>** | $(5.9 \pm 0.8) \cdot 10^{-2}$ | $(1.3 \pm 0.3) \cdot 10^{-4}$ | $(2.0 \pm 0.4) \cdot 10^{-3}$ | $(2.1 \pm 0.7) \cdot 10^{-4}$ |
| **k<sub>cat</sub>/K<sub>M</sub> (wt), M<sup>-1</sup>·s<sup>-1</sup>** | $\sim (1-4) \cdot 10^6$ | $(1.3 \pm 0.1) \cdot 10^4$ | $(3.5 \pm 0.3) \cdot 10^4$ | $(3.4 \pm 0.3) \cdot 10^3$ |
| **k<sub>cat</sub>/K<sub>M</sub> (ha), M<sup>-1</sup>·s<sup>-1</sup>** | $\sim (0.3-2.0) \cdot 10^6$ | $(3.9 \pm 0.4) \cdot 10^2$ | $(5.6 \pm 0.3) \cdot 10^2$ | 69 ± 9 |

### Hyper-accurate RF1_ha_ is dynamic in the A site

To understand the structural basis of the extraordinary termination fidelity of the hyper-accurate Δ302-304 RF1 variant (termed RF1_ha_), we determined crystal structures of 70S•RF1_ha_ termination complexes formed on a UAA stop codon in the absence (Fig. 2A, 2B) and presence of BlaS (Fig. 2C, 2D), at 3.55-Å and 3.4-Å resolutions, respectively (Table 2). We reasoned that destabilizing the GGQ binding at the peptidyl-transferase center by BlaS might increase our chances of capturing an elusive pre-activation conformation of the high-fidelity enzyme. In the absence of BlaS (Fig. 2A), RF1_ha_ adopts an extended (activated) conformation (Fig. 3A) nearly identical to that of RF1wt observed in canonical termination complexes (11, 19). This finding is consistent with our observation that the catalytic activity of RF1_ha_ on the stop codon is similar to that of RF1wt (Fig. 1B-C). The position of the GGQ motif in the peptidyl-transferase center (Fig. 3B) and recognition of the stop codon (Fig. 3C) are similar to those in canonical termination complexes (11, 19), as expected. The interactions of RF1_ha_with the decoding center are destabilized due to shortening of the switch loop (Fig. 3D), however the conformations of decoding center nucleotides A1492, A1493 and A1913 are similar to those in canonical termination complexes.

**Figure 2.**
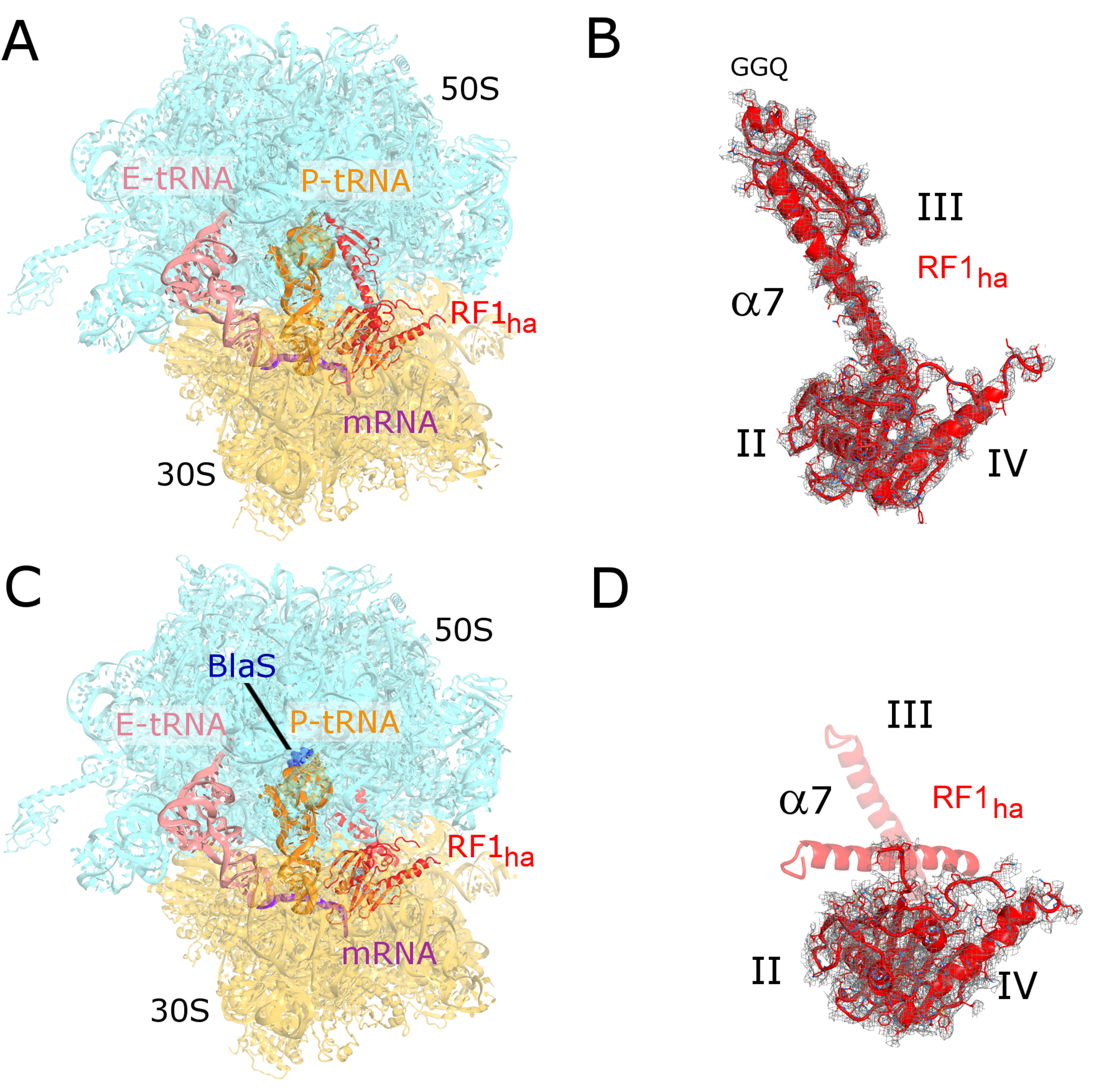
Crystal structures of RF1_ha_ bound to the 70S ribosome in the presence or absence of BlaS. (*A*) Structure of RF1_ha_ bound to the ribosome in the absence of BlaS. RF1_ha_ adopts an extended conformation. (*B*) Close-up view of the release factor and unbiased Fo-Fc densities is shown for RF1_ha_ bound to the ribosome in the absence of BlaS. (*C*) Structure of RF1_ha_ bound to the 70S ribosome in the presence of BlaS. The codon-recognition superdomain (domains II and IV) is bound at the 30S subunit, whereas domain III has partial occupancy in a compact and a partially extended state. (*D*) Close-up view of the release factor and unbiased F_o_-F_c_ densities is shown for RF1_ha_ bound to the ribosome in the presence of BlaS (see Fig. 4 for additional density views).

**Figure 3.**
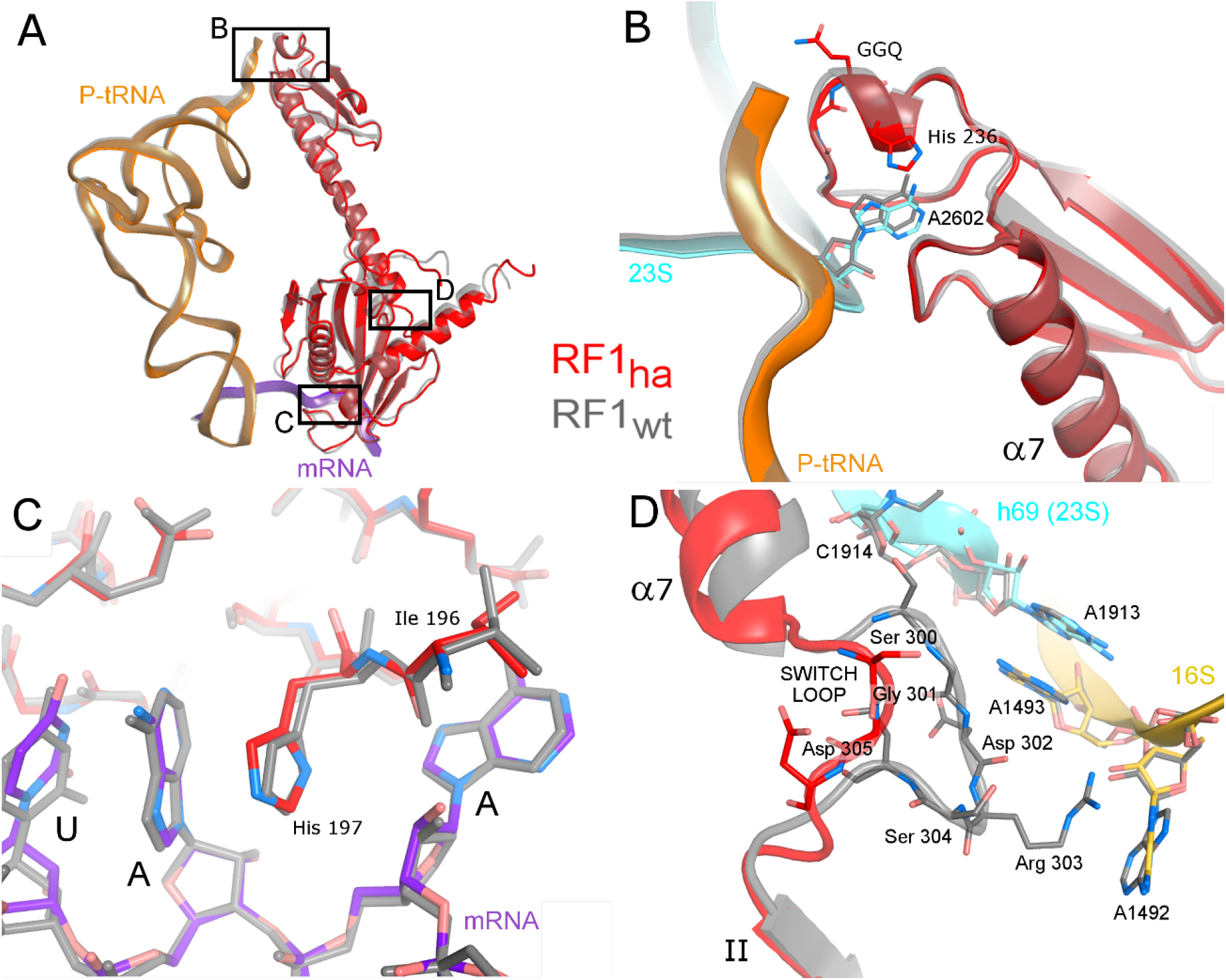
Interactions of RF1_ha_ with the ribosome in the absence of BlaS. (*A*) Alignment of RF1, P-site tRNA and mRNA from the 70S complexes with RF1_ha_ and RF1_wt_ shows that RF1_ha_ adopts the extended conformation characteristic of canonical termination complexes. (*B-C*) Conformation of RF1_ha_ is similar to that of RF1_wt_ in: (*B*) the peptidyl-transferase center and (*C*) the stop-codon-recognition region. (*D*) Positions of the switch loop of RF1_wt_ and RF1_ha_ in the decoding center. The switch loop of wild-type RF1 packs to interact with decoding-center nucleotides, whereas the shortened switch loop of RF1_ha_ does not interact with decoding-center nucleotides. The structure with RF1_ha_ is colored (this work) and the structure with RF1_wt_ is gray (19) (PDB ID 5J4D). The superposition was achieved by structural alignment of 16S rRNAs.

**Table 2.** Data collection and structure refinement statistics.

|  | RF1<sub>ha</sub><br>(PDB **6BOK**) | RF1<sub>ha</sub>-BlaS<br>(PDB **6BOH**) |
| --- | --- | --- |
| **Data collection** |  |  |
| Space group | P2<sub>1</sub>2<sub>1</sub>2<sub>1</sub> | P2<sub>1</sub>2<sub>1</sub>2<sub>1</sub> |
| Cell dimensions |  |  |
| *a*, *b*, *c* (Å) | 210.82, 450.45, 615.17 | 209.86, 450.69, 615.88 |
| $\alpha$, $\beta$, $\gamma$ (°) | 90, 90, 90 | 90, 90, 90 |
| Resolution (Å)* | 70.0 - 3.55 (3.75-3.55) | 70.0 - 3.4 (3.6-3.4) |
| R<sub>p.i.m.</sub><sup>#</sup> | 0.144 (0.982) | 0.129 (1.098) |
| CC (1/2)<sup>##</sup> | 99.7 (34.7) | 99.9 (29.2) |
| I / σI | 7.90 (1.12) | 7.75 (0.96) |
| Completeness (%) | 100 (100) | 100 (100) |
| Redundancy | 38.5 (36.0) | 38.1 (35.3) |
| **Structure Refinement** |  |  |
| Resolution (Å) | 32.4 - 3.55 | 50.0 - 3.4 |
| No. reflections | 697,897 | 792,518 |
| R<sup>work</sup> / R<sup>free</sup> | 0.201 / 0.247 | 0.230 / 0.272 |
| Total No. atoms | 299,596 | 298,216 |
| Ions/water (modeled as Mg<sup>2+</sup>) | 1,518 | 1,428 |
| **R.m.s. deviations** |  |  |
| Bond lengths (Å) | 0.009 | 0.008 |
| Bond angles (°) | 1.238 | 1.197 |
| Coordinate error<sup>†</sup> (Å) | 0.5 | 0.6 |

\* Values in parentheses are for the high-resolution shell.
<sup>#</sup> R<sub>p.i.m.</sub> (precision-indicating merging R factor (46)) was calculated using SCALA, which is part of the CCP4 (47) package.
<sup>##</sup> CC(1/2) is the percentage of correlation between intensities from random half-datasets as defined by Karplus and Diederichs (48).
<sup>†</sup> Maximum-likelihood-based coordinate error is reported (41).

In the presence of BlaS (Fig. 2C, 2D), strong unbiased Fourier difference density reveals that the codon-recognition superdomain of RF1_ha_ (Fig. 4A, 4B) is rotated by ~5° around the stop codon relative to that of wild-type RF1 (Fig. 4D). By contrast, weak density in the intersubunit space suggests partial occupancies for domain III outside of the peptidyl-transferase center (Fig. 4A, 4B), which is bound with BlaS at the same position as in the 70S structure lacking RF1 (29). Although detailed interpretation of these partial occupancies of domain III is difficult due to poor map resolution in this region, it is clear that there is no density corresponding to the docked position of helix α7. Instead, the density is consistent with at least two positions of α7, suggesting a partially extended (mid-open) state of domain III and a compact (closed) domain III (Fig. 4B). The partially extended position of helix α7 deviates from the normally extended conformation by ~20° and is inconsistent with accommodation of the GGQ motif in the peptidyl-transferase center (Fig. 4D). In free release factors, the compact conformation is stabilized by hydrophobic interactions (Fig. 4C) involving O-methyl-serine (Mse) 151, Val 165 and Ile 114 in domain II and Ile 274 of helix α7 in domain III (*Streptococcus mutans* RF1 (24)). These interactions are disrupted in the fully extended release factors (11, 15, (17-20)). In our structure, density in this region (Fig. 4B, 4C) suggests that a fraction of release factors in the crystal may retain some of these interactions, inducing the rotation of the codon-recognition superdomain (Fig. 4D). The steric clash with the P-site tRNA likely forces the GGQ-bearing tip of domain III to displace from its compact conformation, resulting in conformational heterogeneity of domain III. In summary, this structure demonstrates that helix α7 and the GGQ motif are disengaged from the 50S subunit and are destabilized in the intersubunit space.

**Figure 4.**
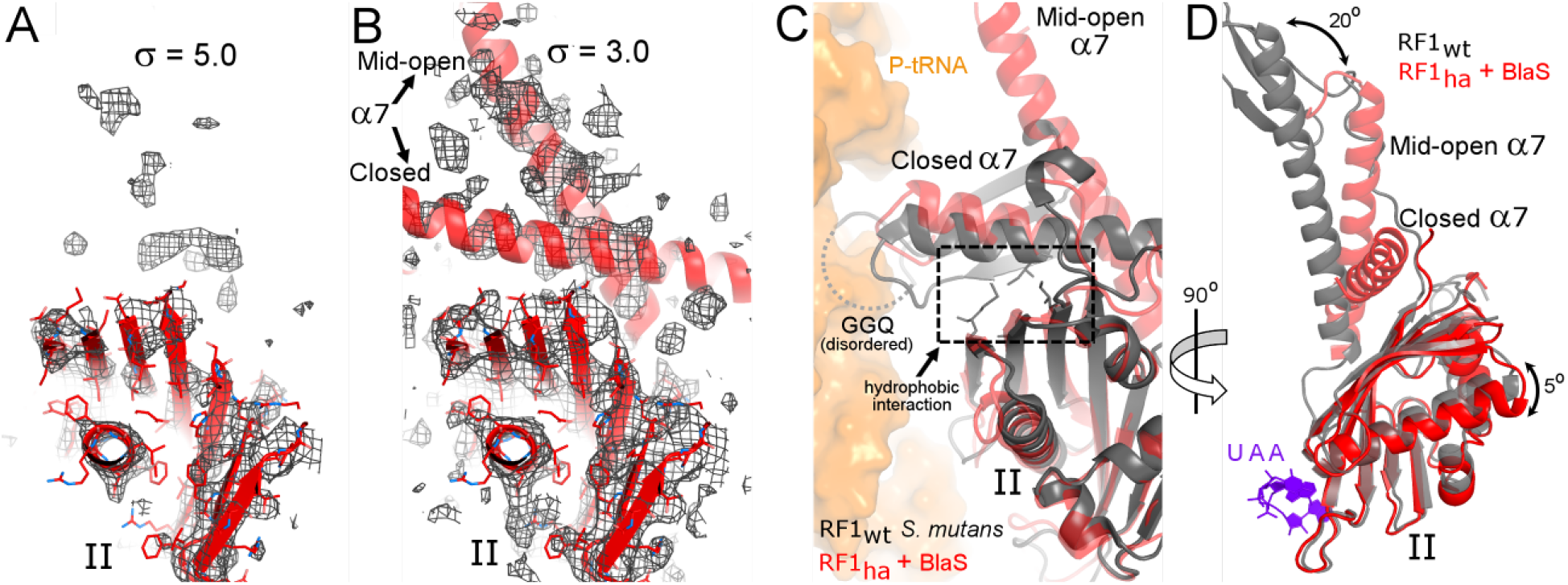
Helix α7 of the catalytic domain III of RF1_ha_ is disengaged from the 50S subunit in the presence of BlaS. (*A*) Unbiased F_o_-F_c_ density (gray mesh) suggests mobility of α7, while the codon-recognition domain II is resolved (at σ=5.0). (B) Low putative occupancies of α7 in the partially extended (mid-open) and compact (closed) conformations are suggested at lower density levels (σ=3.0). (C) Comparison of RF1_ha_ with free (ribosome-unbound) S. *mutans* RF1 (24) (PDB ID 1ZBT; gray) docked into the A site by structural alignment of domains II. Hydrophobic side-chains are shown as sticks. (D) Both conformations are inconsistent with docking of α7 into the peptidyl-transferase center (gray model shows the extended (active) conformation of wild-type RF1 (19) (PDB ID 5J4D).

Despite being bound with the rotated codon-recognition domain and dynamic domain III of RF1 ha, the decoding center of the ribosome adopts the same conformation as in canonical termination complexes. The structure therefore shows that the binding of the codon-recognition domain alone is sufficient to stabilize the termination-specific conformation of the decoding center (Fig. 5).

**Figure 5.**
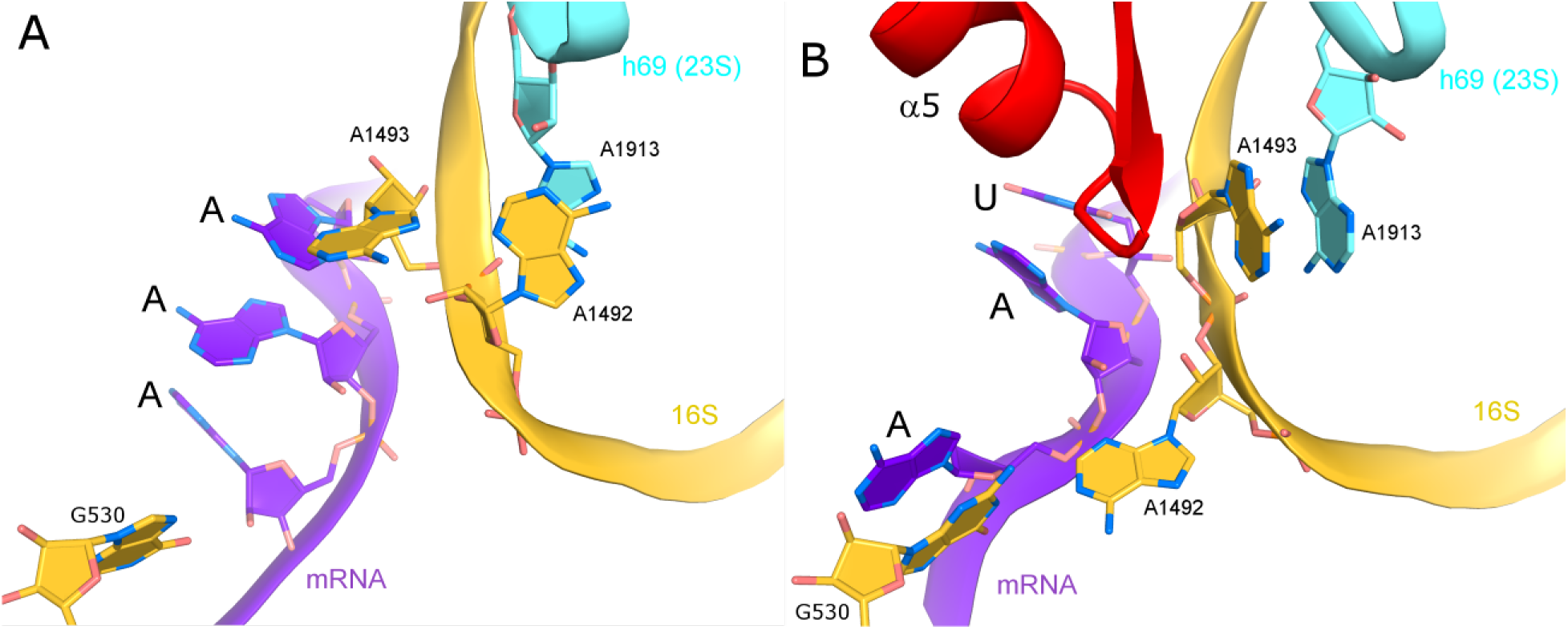
Comparison of the decoding-center nucleotides in the absence *(A; (30))* and presence of RF1_ha_ *(B; without BlaS)*, showing that rearrangement of A1493 is required for the binding of the codon-recognition domain of release factor.

### Conformational control of termination fidelity

New conformations of a release factor captured in our structures provide insight into the mechanism of termination fidelity. They show that the codon-recognition domain can bind the 30S subunit and induce decoding-center rearrangements independently of the switch loop and GGQ accommodation in the peptidyl-transferase center. Moreover, the dynamics of RF1_ha_ are consistent with the hypothesis that release factors undergo a large conformational change upon binding to the stop codon.

Our structures suggest that two principal rearrangements determine the mechanism of termination (Fig. 6): those in the 30S decoding center and those in the release factor. During elongation, release factors sample the ribosome in a somewhat compact conformation (13, 14), perhaps similar to that reported in crystal structures of free release factors (22-24) or compact RF2 bound to the ribosome in the presence of ArfA (27, 28). Prior to RF binding, the decoding-center conformation is incompatible with bound RF. Specifically, crystallographic (30) and cryo-EM (31, 32) structures of the ribosome with a vacant A site show the codon interacting with A1493, which bulges from helix 44 (h44) of 16S rRNA, while A1492 is stacked on A1913 within h44 (Fig. 5A). In this conformation, A1493 would clash with the accommodated RF (Fig. 5B). We propose that transient interaction of RF with a sense codon is insufficient to stabilize the codon-recognition domain on the 30S subunit and induce decoding-center rearrangements compatible with RF binding. In this case, the decoding-center conformation would favor RF dissociation (33) prior to RF domain rearrangement, thus preventing premature termination (Fig. 6B). This idea is consistent with independent proposals that a conformational rearrangement is the rate-limiting step (13, 18, 34). In rare cases, however, RF can induce decoding-center rearrangements on a sense codon and sample the extended state, resulting in premature termination (10) (Fig. 6A).

**Figure 6.**
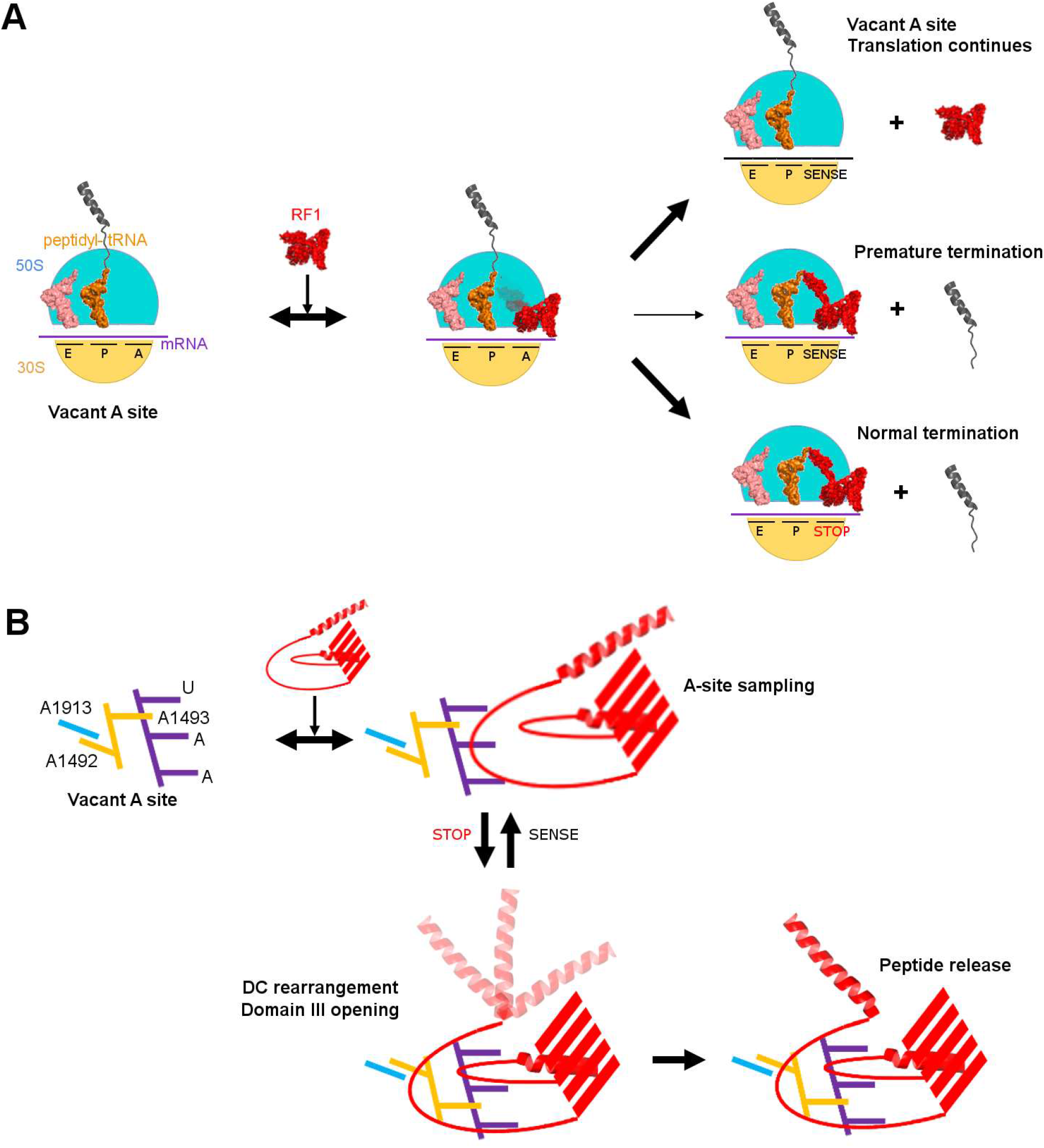
Mechanism of translation termination. (A) Schematic of global conformational rearrangements in release factor. (B) Schematic of local rearrangements in the decoding center (DC) coupled with global rearrangement in the release factor.

When the ribosome reaches the end of the open reading frame, the codon-recognition domain of a release factor stably binds to the 30S A site due to specific interactions with the stop codon. The binding of the codon-recognition domain of RF is coupled with swapping of A1493 with A1492, so that A1493 stacks on A1913 and A1492 bulges out of h44 (Fig. 6B). Our structures demonstrate that interaction of the switch loop or the extended domain III with the decoding center are not required for this decoding-center rearrangement. Thus, the binding of the codon-recognition domain to the stop codon stabilizes the rearranged decoding-center nucleotides prior to the activation of RF.

Stable binding to the stop codon provides an opportunity for domain III to disengage from the codon-recognition superdomain and sample different conformations in the intersubunit space. The loss of hydrophobic interactions between domains II and III is compensated by new interactions between the switch loop and the decoding center, and between the extended domain III and the large subunit. The insertion of the GGQ motif into the peptidyl-transferase center is coupled with changes in peptidyl-transferase-center nucleotide conformations, as described earlier (reviewed in (1)). We have recently determined the structure of the 70S complex formed with wild-type RF1 and BlaS (35). The structure revealed that BlaS perturbs the peptidyl-transferase center and shifts the GGQ motif of RF1 without destabilizing the catalytic domain overall. Our current work demonstrates that perturbation of the switch loop by mutation shifts the equilibrium away from the extended conformation. Thus, the wild-type switch loop plays a critical role in stabilizing the extended active RF1 conformation in the ribosome. Our biochemical data suggest that stabilization of the extended RF1 conformation reduces termination fidelity but likely provides a kinetic advantage under cellular conditions, thereby balancing the accuracy of termination, the optimal translation rate and ribosome recycling.

## Acknowledgements

We thank Rohini Madireddy and Gabriel Demo for help with ribosome and protein purification, staff of Argonne National Laboratory (Advanced Photon Source, beam lines 23 ID-B, 23 ID-D), SLAC National Accelerator Laboratory (Stanford Synchrotron Radiation Lightsource, beam lines 9-2, 12-2), and Brookhaven National Laboratory (National Synchrotron Light Source I, beam line X25) for the assistance with data collection; Emily Mohn and Darryl Conte Jr. for the assistance with manuscript preparation; Dmitri Ermolenko and Korostelev lab members for discussions and comments on the manuscript. This study was supported by NIH grants R01 GM107465 and P30 DK047757 (to A.A.K.). X-ray structures have been deposited to RCSB (PDB 6BOH and 6BOK).

## Materials and Methods

### Purification of Ribosomes and Release Factors

*Escherichia coli* 70S ribosomes used in biochemical analyses were purified from MRE600 cells (36). *Thermus thermophilus* ribosomes (HB 27) and *E. coli* RF1_ha_ (Δ302-304 RF1) were used for crystallization. C-terminally His-tagged *E. coli* RF1_ha_ and *E. coli* RF1_wt_, used for peptide release assays, were purified as described (11). We previously demonstrated that *E. coli* RF1 is active on *T. thermophilus* 70S ribosomes and that this heterologous system is suitable for crystallographic studies of termination (19).

### Peptide Release Assays

*E. coli* tRNA^fMet^ (Chemical Block) was aminoacylated using [^35^S]-methionine (Perkin Elmer) as previously described (37). Peptide release assays were performed as described (19). The pre-termination complex was prepared by mixing [^35^S]-fMet-tRNA^fMet^ with 1.5-fold molar excess of *E. coli* 70S ribosomes and incubating for 15 minutes at 37°C in buffer A (20 mM Tris·HCl (pH 7.5), 100 mM ammonium acetate, 20 mM magnesium acetate). After incubation, we diluted the complex ~20-fold to 10 nM ([^35^S]-fMet-tRNA^fMet^) in buffer A supplemented with 0.05% Triton X-100 (OmniPur) and a messenger RNA GGCAAGGAGGUAAAA<u>AUG**XYZ**</u>AAAAAA (XYZ stands for the A-site codon following the AUG codon placed in the P site; IDT) at 1 μM (final concentration). After incubation for 5 minutes at 37°C, the pre-termination complex was cooled to room temperature.

In the kinetic assay, an aliquot (4.5 μl) of the pre-termination complex was quenched in 30 μl of 0.1 M HCl to represent the zero-time point. 45 μl of the pre-termination complex were combined with 5 μl of a 10-fold release factor solution. 5 μl aliquots were quenched at different time points in 30 μl of 0.1 M HCl. Every quenched aliquot was extracted with 700 μl of ethylacetate and 600 μl of the extract were mixed with 3.5 ml of scintillation liquid (Econo-Safe). The amount of released [^35^S]-labeled N-formyl-methionine in each aliquot was determined using a scintillation counter (Beckman Coulter, Inc.). Fitting time progress curves to a single exponential function yielded rate constants (k_obs_). RF1 concentrations and k_obs_ values were fitted to a hyperbola in order to obtain apparent k_cat_ values. Enzyme efficiencies (k_cat_/K_M_) were determined using a linear approximation of the Michaelis – Menten equation, omitting data points equal or greater than estimated KM values. GraphPad Prism 7 was used for data plotting and graph drawing. Each time-progress-curve data point was obtained from two independent experiments.

### Crystallization, Data Collection and Processing

Crystallization of 70S•RF1 complexes formed with the UAA-containing mRNA (GGCAAGGAGGUAAAA<u>AUG</u>**UAA**AAAAAA) was performed essentially as described (19, 29, 38). 4 μM (all concentrations are given for the final crystallization solution) *T. thermophilus* ribosome was incubated with 12 μM UAA-containing mRNA, 10 μM tRNA^fMet^, and 16 μM *E. coli* RF1 (RF1_wt_ or RF1_ha_) in the buffer containing 25 mM Tris acetate (pH 7.0), 50 mM potassium acetate, 10 mM ammonium acetate, 10 mM magnesium acetate and 2.8 mM Deoxy Big CHAPS (Soltec Ventures). 1 mM BlaS (Fischer Scientific) was supplemented to this solution to form 70S•RF1•BlaS complex. Crystals were obtained using hanging-drop vapor diffusion over 300 μl of 0.4-0.6 M NaCl by combining 3 μl of the ribosome complex and 3 μl of crystallization solution containing 0.1 M Tris·HCl (pH 7.5), 4% (v/v) PEG 20000 (Hampton Research), 8% (v/v) 2-Methyl-2,4-pentanediol (Hampton Research), and 0.2 M KSCN. Crystals were cryo-protected in four overnight steps, in which the composition of the mother liquor was incrementally adjusted to 0.1 M Tris·HCl (pH 7.5), 4.5% (v/v) PEG 20000, 10% (v/v) PEG 200 (Hampton Research), 30% (v/v) 2-methyl-2,4-pentanediol and 0.2 M KSCN. During cryo-protection of complex that contained BlaS, all cryo-protection solutions were supplemented with 0.5 mM BlaS. Crystals were frozen by plunging into liquid nitrogen.

Crystals were screened at Argonne National Laboratory (Advanced Photon Source, beam lines 23 ID-B, 23 ID-D), SLAC National Accelerator Laboratory (Stanford Synchrotron Radiation Lightsource, beam lines 9-2, 12-2), and Brookhaven National Laboratory (National Synchrotron Light Source I, beam line X25). The data that were used for structure determination were obtained at Argonne National Lab (Advanced Photon Source, beam line 23 ID-D, 1.033 Å wavelength, detector Dectris PILATUS3 6M) and Brookhaven National Laboratory (National Synchrotron Light Source I, beam line X25, 1.1 Å wavelength, detector PILATUS 6M PAD) using the 0.2° oscillation range.

The final data set for each structure was obtained by merging six data sets for the complex with RF1_ha_, and five data sets for the complex with RF1_ha_ and BlaS. The data were scaled with XDS (39). Two percent of reflections in each dataset were used as a test set (R^free^ set). The starting model for molecular replacement was obtained from the previously reported structure with *E. coli* RF1 (PDB ID: 5J4D (19)), from which the stop codon, RF1 (for all complexes), and nucleotides 74-76 (CCA) of P-tRNA (for complexes with BlaS) were removed. Densities for the stop codon, RF1 codon-recognition superdomain, CCA end of the P-site tRNA and BlaS were clearly visible in unbiased F_o_-F_c_ and FEM maps (40) calculated using PHENIX (41). We found BlaS in the same position as previously described (29). The distorted CCA end of the P-site tRNA with the intercalated BlaS in the presence of RF1 adopts the same conformation as in the complex lacking RF1 (29). Density for domain I of RF1 in the vicinity of the L11 stalk was weak in all structures reported here, as in the previously reported crystal structure of the 70S•RF1 complex (11), consistent with dynamic domain I also observed in crystal structures of *Thermus thermophilus* 70S•RF1 complexes (11, 19). Magnesium ions were modeled using CNS (42, 43). NCS-restrained simulated-annealing refinement in combination with group B-factor and TLS refinements was carried out using PHENIX (41). Data and refinement statistics are summarized in the Table 1. Figures were rendered using PyMOL (44). Rotation angles and distances between atoms were measured using Chimera (45) and PyMOL, respectively.

